# The Joubert gene *TMEM67* is required for the correct establishment of spinal dorsal identities in human organoids

**DOI:** 10.64898/2026.01.20.700675

**Authors:** A. Wiegering, K. Bools, I. Anselme, L. Metayer-Derout, O. Mercey, E. Balissat, Y. Bijek, M. Catala, S. Schneider-Maunoury, A. Stedman

## Abstract

Primary cilia are essential signaling organelles that mediate key developmental pathways, including Sonic Hedgehog (SHH) and WNT, and are crucial for tissue patterning and homeostasis. Ciliary dysfunction underlies a spectrum of human ciliopathies—such as Joubert syndrome (JBTS) and Meckel syndrome (MKS)—which present with profound neurodevelopmental abnormalities. Although the role of cilia in SHH-dependent ventral spinal cord patterning is well established, their contribution to dorsal spinal cord development, particularly in human systems, remains poorly defined. To address this gap, we utilized human spinal organoids to investigate the function of a ciliopathy-associated protein in dorsal neural tube patterning downstream of BMP4, independent of exogenous SHH. Using *TMEM67* knockout human iPSC-derived dorsal spinal organoids, we demonstrate that loss of TMEM67 disrupts the specification of dorsal interneurons, most notably within the dI1 lineage, while concomitantly expanding intermediate dorsal progenitor populations (dI4–dI6). These patterning defects are associated with defective roof plate induction and attenuated BMP4 signaling. Mechanistically, TMEM67 deficiency alters ciliary morphology, decreases cilia number, and impairs recruitment of BMPR2 to the ciliary base, suggesting a direct role for cilia in modulating BMP-induced dorsal spinal patterning. Together, these findings provide new mechanistic insights into the pathogenesis of ciliopathies and underscore the value of human organoid models for elucidating human-specific aspects of neurodevelopmental disorders.

## INTRODUCTION

Primary cilia are sensory and signalling organelles present on virtually every cell during development and in adults. They transduce signalling pathways essential for development and homeostasis, such as Hedgehog (HH), Wnt and many others (reviewed in^1^). Cilia are enriched in signal receptors and transducers thanks to the transition zone, a proximal region of the ciliary axoneme involved in controlling protein trafficking to and from the cilium. In line with these essential signaling functions, ddysfunction of primary cilia lead to ciliopathies, a clinically and genetically heterogenous group of diseases that can affect multiple organs such as the kidney, retina, liver, skeleton and brain (reviewed in^2, 3^). Joubert syndrome (JBTS) and Meckel syndrome (MKS) are two neurodevelopmental ciliopathies with distinct severities. Meckel syndrome is characterized by occipital encephalocele and severe brain morphological defects and is lethal at birth, while the main feature of JBTS is brainstem and cerebellar dysgenesis, with variable forebrain anomalies (for review^4^). Accordingly, mouse mutants of JBTS- and MKS-causal genes display multiple brain defects ^5, 6, 7, 8, 9, 10, 11, 12, 13^. The complexity and intricacy of these defects claim for the use of simpler systems to study the role of cilia in signalling and neural development.

Over the years, the developing spinal cord has become a paradigm for the study of signalling pathways in neural tube patterning (reviewed in^14^). Using this paradigm, seminal studies in mice have uncovered the crucial role of primary cilia in Hedgehog/Gli (Hh) signalling and cell type specification^15^, reviewed in^16^. The developing spinal cord is patterned along its dorso- ventral (DV) axis by to the coordinated activity of four main signalling pathways: Sonic Hedgehog (SHH) ventrally, BMPs and WNTs dorsally and retinoic acid (RA) laterally. Sonic Hedgehog acts as a morphogen, specifying different neural progenitor types along the ventral spinal cord depending on their distance from the morphogen source and time of exposure^14^. BMPs and WNTs expressed by the roof plate are necessary for the specification of the most dorsal cell types^17, 18, 19^, reviewed in^20^. Cilia-less mutants show a loss or strong reduction of ventral cell types, from floor plate cells to motoneurons, due to a loss of the activator form of the Gli2 transcription factor^15^. Functional cilia are also required for the formation of the repressor form of Gli3, normally found in cells receiving little or no HH signals. In consequence, cells totally devoid of cilia show a basal level of Gli transcription factor activity, with little activation but also little repression of the pathway (reviewed in^16, 4^).

In mouse mutants for individual ciliopathy genes, perturbation of Gli activity and spinal cord patterning are variable, as exemplified for *TMEM67, RPGRIP1L* and *ARL13B*, three JBTS- and MKS- causal genes^21, 22, 23^. In the *Rpgrip1l* mutant mouse (also called *Ftm*), the floor plate and the V3 interneurons, which require high levels of Hh signalling for their production, are absent, while motoneurons, which require intermediate levels of Hh signalling, are reduced in number^24^. In the *Arl13b/Hnn* mutant, the floor plate is absent but motoneurons are increased in number and their domain is expanded both dorsally and ventrally, due to a shallow gradient of Hh/Gli activity^25^. In *Tmem67* mutants, both situations can occur depending on the genetic background^8^.

The role of primary cilia in the dorsal spinal cord has been much less studied. In the *Arl13b/ Hnn* mouse mutant, the most dorsal spinal cord cell types in the lumbar region are strongly reduced, with loss of Gdf7 (a marker of the roof plate) and of the dorsal markers Msx1 and Lhx2. Intriguingly, this phenotype is not present in the more anterior spinal cord, in which dorsal cell populations form normally. Moreover, this phenotype is not reproduced when *Arl13b* is conditionally inactivated in the dorsal Pax3-positive progenitor domain from E10.5 onward, leading the authors to conclude that the *Hnn* phenotype is and indirect cause of the ectopic activation of Shh/Gli signalling in the dorsal spinal cord^26^. This conclusion is comforted by the analysis of mouse mutants for other ciliary genes that display over-activated HH signalling, such as *Fkbp8* and *Tulp3*, which show a reduction of dorsal spinal cord cell types^27, 28^.

Whether defects in dorsal spinal cord cell fate specification can arise independently of HH/Gli deregulation in ciliary mutants remains to be determined. Moreover, the spinal cord has been little examined in JBTS patients, despite a couple of papers that describe histological anomalies of the dorsal horns^29, 30^, and even less so in Meckel fetuses, for which very few samples are available. Thus, little is known about whether and how cilia may affect human dorsal spinal cord cell fates. To address these questions, we turned to human iPSC-derived spinal organoids. By recapitulating developmental pathways, these approaches are well suited to study early steps of neural tube development and patterning^31, 32^. Using ventral spinal differentiation from mouse ESCs and human iPSCs, we recently identified significant differences in cilia formation and function in human and mouse spinal organoids in the absence of the JBTS proteins RPGRIP1L and TMEM67^33^. This further emphasizes the importance of using human-based systems to investigate the developmental origin of cilia-related neural defect.

In this paper we generate dorsal spinal organoids from human iPSCs KO for the JBTS gene *TMEM67* alongside isogenic controls. *TMEM67* encodes a frizzled-type transmembrane protein localized at the transition zone. Combining bulk RNAseq, qPCR and immunofluorescence, we show that *TMEM67* KO human spinal dorsal organoids exhibit disrupted specification of the most dorsal interneuron subtypes, while intermediate dorsal lineages are expanded or unaffected. These patterning defects coincide with impaired roof plate formation and attenuated response to BMP4 signalling. Finally, we uncover a recruitment of BMPR2 at the ciliary base in dorsal spinal progenitor cells, which is reduced in absence of TMEM67. Together, our findings reveal a previously unrecognized and direct role for primary cilia and ciliopathy genes in dorsal neural patterning.

## RESULTS

### TMEM67 loss is associated with reduced dl1–dl3 markers and increased dl4–dl6 neuronal identities in human dorsal spinal organoids

The dorsal neural tube forms six neural progenitor types, dp1 to dp6, as well as a group of very dorsal cells that gives rise to neural crest cells (NCCs) and the roof plate (RP) (Figure 1A). dp1 to dp3 generate dI1 to dI3 relay interneuron (IN) subtypes, while dp4 to dp6 produce five associating IN subtypes, first dI4 to dI6 and later dILA and dILB. To investigate the role of the ciliopathy gene *TMEM67* in human dorsal spinal cord patterning, we generated dorsal spinal organoids from a heterozygous control and a *TMEM67* homozygous knockout hiPSCs clones (TMEM67^+/-^ and *TMEM67*^-/-^, hereafter called *CTRL* and *TMEM67^KO^*) using a previously published protocol^32^ (Figure 1B). In this system, BMP4 treatment of hiPSC-derived embryoid bodies robustly induces roof plate and dl1–dl6 dorsal identities in a highly reproducible spatiotemporal sequence. This process results in a concentric organization of dorsal progenitor domains, with progressively more dorsal identities positioned toward the outer layers, faithfully recapitulating the stepwise dorsal regionalization of the neural tube *in vivo*.

**Figure 1:**
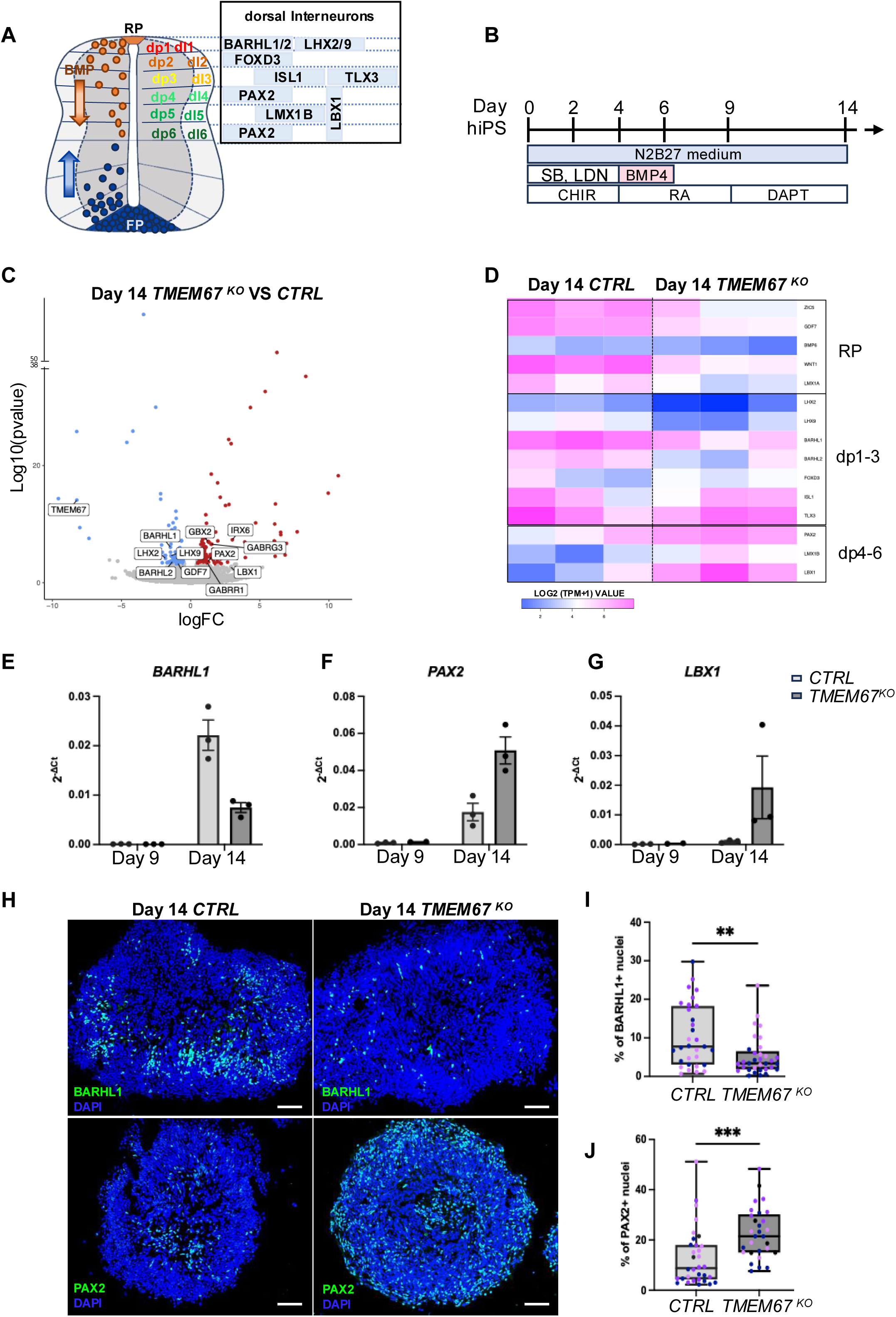
*TMEM67*-deficient dorsal spinal organoids fail to adopt dorsal-most, BMP-dependent neuronal fates. **(A)** Diagram depicting the main transcription factors spanning the different neuronal domains of the developing dorsal spinal cord. **(B)** Schematic summary of the spinal 3D differentiation protocol. Small molecules used to induce or repress signaling pathways are indicated below. SB and LDN: SMAD antagonists SB-431542 and LDN-193189; CHIR: WNT agonist CHIR99021; BMP4: recombinant Bone Morphogenetic Protein 4; DAPT: NOTCH agonist; RA: Retinoic Acid. **(C)** Volcano Plot showing DEGs between *TMEM67^KO^* and *CTRL* organoids at day 14, obtained from bulk RNAseq, N=3 experimental replicates per genotype (genes with P<0.05; Log2FC>1 are colored). **(D)** Heatmap visualization of normalized expression (log2(tpm+1) values) of a distinctive set of dorsal neuronal markers in *TMEM67 ^KO^* and *CTRL* organoids at day 14. Each column represents an experimental replicate. **(E-G)** qRT-PCR for indicated genes in *CTRL* and *TMEM67^KO^* and organoids at day 9 and day 14. N=3 experimental replicate per genotype. Each dot represents a distinct experimental replicate. Data shown are mean ± SEM. **(H)** Immunofluorescence for BARHL1 (dI1) and PAX2 (dI4 and dI6) in sections from *TMEM67^KO^* and *CTRL* organoids at day 14. **(I, J)** Quantification of the percentage of stained nuclei for the indicated markers in sections from *CTRL* and *TMEM67^KO^*organoids at day 14. N=3 experimental replicates per genotype. Each color represents a distinct experimental replicate. Data are shown as mean ± SEM. **P < 0.05, ***P < 0.001 (unpaired t tests with Welch’s correction). Scale bar, 50 µm.

Organoids from 3 independent differentiation experiments performed with 1 clone of each genotype were collected on days 4, 9, and 14 of differentiation and analysed by bulk RNAseq. These stages correspond respectively to: day 4, when organoids are predominantly composed of axial neuro-mesodermal progenitors; day 9, when spinal neural progenitors are specified and day 14, marking the onset of spinal interneuron differentiation and maturation. To verify that both control and mutant organoids were able to differentiate into dorsal spinal organoids, we first assessed their global transcriptional trajectories. Principal component analysis (PCA) showed that samples segregated primarily along PC1 and PC2 according to differentiation time, before separating by genotype (Supplementary Figure 1A). This indicates that temporal progression of differentiation is the dominant source of transcriptional variation in both conditions. Consistently, analysis of stage-specific markers revealed appropriate and comparable differentiation dynamics in control and TMEM67-deficient organoids, confirming that the overall progression of spinal cord differentiation followed the anticipated temporal program dictated by the protocol (Supplementary Figure 1B). Importantly, analysis of the differentially expressed genes at day 14 indicated a ventral shift of *TMEM67 ^KO^* dorsal spinal organoids relative to controls. As shown in the volcano plot (Figure 1C), *GDF7*, expressed in RP cells, and several hallmark dI1 transcription factors—including *BARHL1*, *BARHL2*, *LHX2*, and *LHX9*—were significantly downregulated in *TMEM67*-deficient organoids. In contrast, *PAX2*, a marker of dl4 and dl6 interneurons, was markedly upregulated. In line with this shift, genes associated with GABAergic inhibitory neuron identity—characteristic of dl4–dl6 populations—such as *GABRG3* and *GABRR1*, were upregulated in mutant organoids. Based on normalized expression of dorsal spinal markers visualized as a heatmap, we observed that at day 14 of the differentiation, the molecular profile of post-mitotic interneurons shifted from RP/dl1–dl3 identities in control organoids to dl4–dl6 identities in the mutant (Figure 1D). Changes in the expression of *BARHL1* (dl1), *PAX2* (dl4/6) and *LBX1* (dl4-dl6) were consistently confirmed by RT-qPCR across three independent differentiations (Figure 1E-1G). To validate these findings at the protein level—and leveraging the stereotyped spatial organization of nuclei expressing dorsal markers, which form concentric layers mirroring dorso–ventral patterning in organoids—we performed immunofluorescence for BARHL1 (dl1) and PAX2 (dl4/dl6). In line with our molecular analysis, we observed a marked reduction in the number of BARHL1-positive neurons (Figure 1H, 1I) accompanied by an increase in PAX2- positive neurons (Figure 1H, 1J) in *TMEM67^KO^*organoids compared to controls. Notably, the spatial distribution of PAX2-positive cells differed between control and *TMEM67^KO^* organoids. In mutant organoids, these cells were enriched in the outermost layers—regions typically occupied by more dorsally specified cells—further supporting altered dorsal patterning of spinal progenitors in absence of TMEM67.

### TMEM67 deficiency is associated with altered roof plate and dp1 specification and reduced BMP signaling

To determine whether the reduction in RP cells and dI1–3 interneurons observed in the absence of TMEM67 arises from early defects in dorsal patterning, we analyzed the expression of markers characteristic of distinct dorsal progenitor domains (Figure 2A). Differential expression analysis revealed a significant reduction in the dp1 lineage marker *ATOH1*, as illustrated in the volcano plot (Figure 2B). This result was further validated by immunostaining (Figure 2C, D) and qRT-PCR (Figure 2E), which revealed an even more pronounced decrease in *ATOH1*-expressing progenitors by day 14. Together, these findings suggest that the reduced dl1 marker expression observed at later stages likely arises from early defects in dp1 lineage induction. Although not statistically significant, *OLIG3* expression—which labels the dp1–dp3 domain—was moderately decreased at both the mRNA and protein levels and displayed a patchier staining pattern in mutant organoids (Figure 2G, H-I), likely reflecting a less penetrant dysregulation of the p2/3 domain compared with p1. In contrast, as expected given their broad expression across the developing dorsal spinal cord, *PAX3* and PAX7 (dp1–dp6) levels were unchanged (Figure2H, J, K and Supplementary Fig. 1C–E).

**Figure 2:**
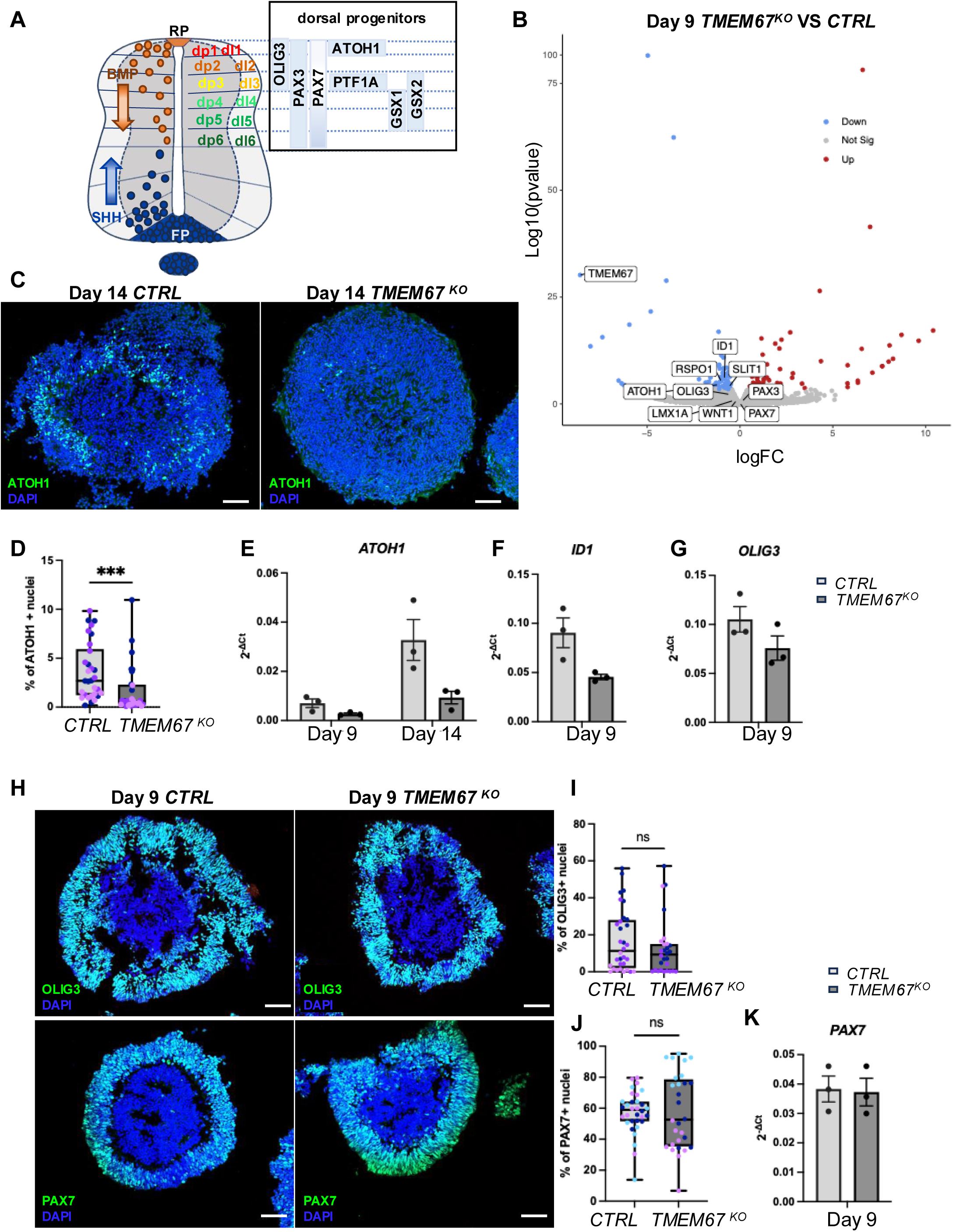
*TMEM67*-deficient dorsal spinal organoids show early defects in the specification of dorsal identities downstream of BMP4. **(A)** Diagram depicting the main transcription factors spanning the different progenitor domains of the developing dorsal spinal cord. **(B)** Volcano Plot showing DEGs between *CTRL* and *TMEM67 ^KO^* organoids at day 9, obtained from bulk RNAseq, N=3 experimental replicates per genotype (genes with P<0.05; Log2FC>1 are colored). **(C)** Immunofluorescence for ATOH1 in sections from *CTRL* and *TMEM67^KO^* organoids at day 14. **(D)** Quantification of the percentage of ATOH1-positive nuclei in sections from *CTRL* and *TMEM67^KO^* organoids at day 14. N=3 experimental replicates per genotype. Each color represents a distinct experimental replicate. Data are shown as mean ± SEM. ***P < 0.001 (unpaired t tests with Welch’s correction). **(E-G)** qRT-PCR for indicated genes in *CTRL* and *TMEM67 ^KO^* organoids at indicated time points. N=3 experimental replicate per genotype. Each dot represents a distinct experimental replicate. Data shown are mean ± SEM. **(H)** Immunofluorescence for OLIG3 and PAX7 in sections from *CTRL* and *TMEM67^KO^* organoids at day 9. **(I, J)** Quantification of the percentage of stained nuclei for the indicated markers in sections from *CTRL* and *TMEM67 ^KO^* organoids at day 9. N=3 experimental replicates per genotype. Each color represents a distinct experimental replicate. Data are shown as mean ± SEM. (unpaired t tests with Welch’s correction). **(K)** qRT-PCR for *PAX7* in *CTRL* and *TMEM67^KO^* organoids at day 9. N=3 experimental replicates per genotype. Each dot represents a distinct experimental replicate. Data shown are mean ± SEM. Scale bar, 50 µm.

Altogether, these phenotypes closely resemble those reported in mouse mutants with impaired roof-plate-derived BMP or WNT signaling. Similar defects have been described in Bmpr1a/b mutants, in which spinal progenitors exhibit reduced responsiveness to BMP cues, leading to a loss of dp1 progenitors accompanied by a dorsal expansion of more ventral progenitor domains^19^. Defects in the specification of dI1 interneurons have also been observed upon decrease of GDF7 or in the Dreher mouse mutant^34, 35^. In addition, compound Wnt1/Wnt3 mutants display a similar reduction in the dp1 progenitor domain^18^. The altered dorsal patterning observed in *TMEM67^KO^*organoids may therefore result from incomplete roof plate induction and/or impaired transduction of roof-plate-derived signals. Consistent with both possibilities, *RSPO1* and *SLIT1*—markers enriched in the early roof plate—were significantly downregulated in *TMEM67^KO^* organoids at day 9 (Figure 2B). Moreover, expression of *ID1*, a direct transcriptional target of roof-plate-derived BMP signaling, was markedly reduced in *TMEM67^KO^* organoids compared with controls (Figure 2B, F). Since roof plate induction and signaling activity in our dorsal organoid differentiation protocol rely on the addition of exogenous BMP4, we first examined the direct cellular response to BMP4 in control and mutant organoids. To this end, control and *TMEM67^KO^* organoids were exposed to 5 ng/ml BMP4 for 1 hour and subsequently immunostained for phosphorylated SMAD1/5/9. Consistent with previous reports^32, 36^, BMP4 treatment induced robust nuclear accumulation of phospho-

SMAD1/5/9 in cells located at the outermost layer of the organoids (Figure 3A, B). While mutant progenitors at the organoid surface were capable of activating BMP signaling similarly to controls, the mean phospho-SMAD–positive area was significantly reduced in *TMEM67^KO^* organoids, suggesting an attenuated or delayed response to BMP4 signaling (Figure 3A-C). To determine whether this reduced response resulted from altered expression of core components of the BMP pathway, we analyzed our bulk RNA-seq data. We found no significant differences between control and *TMEM67^KO^*organoids in the expression of major BMP receptors, SMAD proteins, or known BMP inhibitors at this developmental stage (Figure 3D-G, Supplementary Figure 2A–C). These results indicate that the diminished BMP response observed in *TMEM67^KO^* organoids is unlikely to arise from defects in pathway component expression, but rather from impaired signal transduction. We next investigated activation of the WNT pathway. In particular, we examined the expression of the Wnt ligands WNT1 and WNT3A, which act downstream of BMP signaling to promote the proliferation and maintenance of dorsal progenitors, as well as AXIN2 as a readout of canonical Wnt pathway activity. No significant differences in the expression of these genes were detected between control and mutant organoids (Supplementary Figure 3D–F).

**Figure 3:**
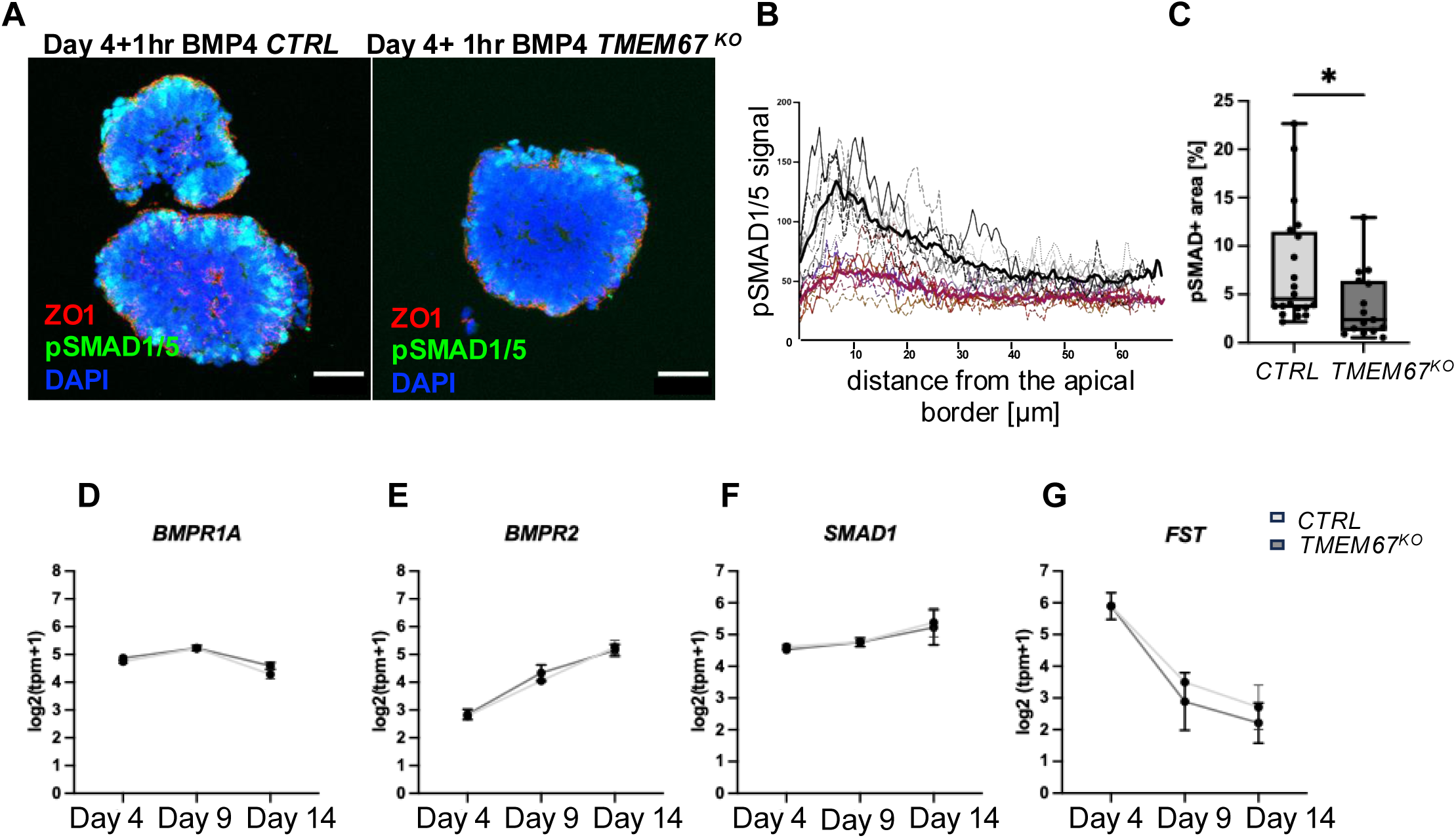
Weakened BMP4 signaling in TMEM67-deficient dorsal spinal organoids. **(A)** Immunofluorescence for P-SMAD1/5/9 on sections of *CTRL* and *TMEM67^KO^* spinal organoids at day 4, 1 hour post BMP4 treatment. Apical cell borders are labeled by ZO1 in red. P-SMAD1/5/9 relative to the distance from the apical border of the organoids stained by ZO1. **(C)** Quantification of the P-SMAD1/5/9-stained area per organoid section. N=1 experimental replicate per genotype. Each dot represents a distinct organoid. Data are shown as mean ± SEM. *P < 0.05. **(D-G)** normalized expression (Log2tpm+1) for indicated genes in *CTRL* and *TMEM67 ^KO^* organoids at indicated time points. N=3 experimental replicate per genotype. Data shown are mean± SEM.

Finally, since SHH and BMP exert opposing roles along the dorsal-ventral axis and ectopic activation of the SHH pathway has been proposed to antagonize BMP-mediated dorsalization in mouse ciliary mutants^26, 37^, we assessed the expression of SHH and its downstream GLI effectors. While SHH expression was barely detectable in both conditions, we observed an upregulation of *GLI1* and *PTCH1* at day 9 in *TMEM67^KO^* organoids (Supplementary Figure 3G–K), suggesting a subtle increase in SHH pathway activity that may contribute to the impaired BMP-dependent dorsalization observed in mutant organoids.

### Loss of TMEM67 in human spinal progenitors leads to defective ciliary gating and altered ciliary localization of BMPR2

Across different cellular contexts, loss of TMEM67 was shown to lead to impaired primary cilia, with variable effects on ciliogenesis, cilium length, and morphology. Despite this variability, impaired ciliary signaling emerge as a common feature, characterized notably by decreased ARL13B recruitment at the ciliary membrane^38, 39, 9^. Given that primary cilia serve as key signaling hubs, and that several BMP pathway effectors have been reported to localize to the ciliary axoneme or basal body in different contexts—sometimes playing a direct role in pathway activation^40, 9^—we next analyzed the ciliary phenotype of progenitors at day 4 following BMP4 stimulation and at day 9.

We found that loss of TMEM67 resulted in a significant reduction in cilia number in organoids at day 4 and 9 of the differentiation (Figure 4A, B, E, H). In addition, TMEM67-associated ciliary phenotypes varied across developmental stages. Although ciliary length was reduced in mutant organoids at day 4 (Figure 4A, C) it was increased at day 9 in dorsal spinal progenitors (Figure 4A, D). We next assessed the gating function of TMEM67 by examining ARL13B recruitment to the ciliary membrane. Throughout dorsal spinal differentiation, TMEM67- deficient progenitors exhibited a marked reduction in ARL13B ciliary localization, consistent with a defect in ciliary gating in *TMEM67^KO^*progenitors (Figure 4A, B, F, G). We used Ultrastructure Expansion Microscopy (U-ExM)^41^ to enhance the resolution of ciliary mapping and to assess the subcellular localization of BMP pathway effectors. Although SMAD1 and phosphorylated SMAD1/5/9 could not be detected at the cilium (data not shown), we observed a striking ciliary localization of BMPR2. BMPR2 localized to multiple ciliary compartments but was particularly enriched at the basal body (Figure 4I). Notably, this basal body enrichment was significantly reduced in *TMEM67^KO^* progenitors (Figure 4I, J). Altogether, these results indicate that TMEM67 is required for proper localization of the BMPR2 receptor at the ciliary base and suggest that, together with additional yet-unknown mechanisms, this defect may contribute to the attenuated BMP4 responsiveness of mutant spinal progenitors, consequently altering dorsal patterning.

**Figure 4:**
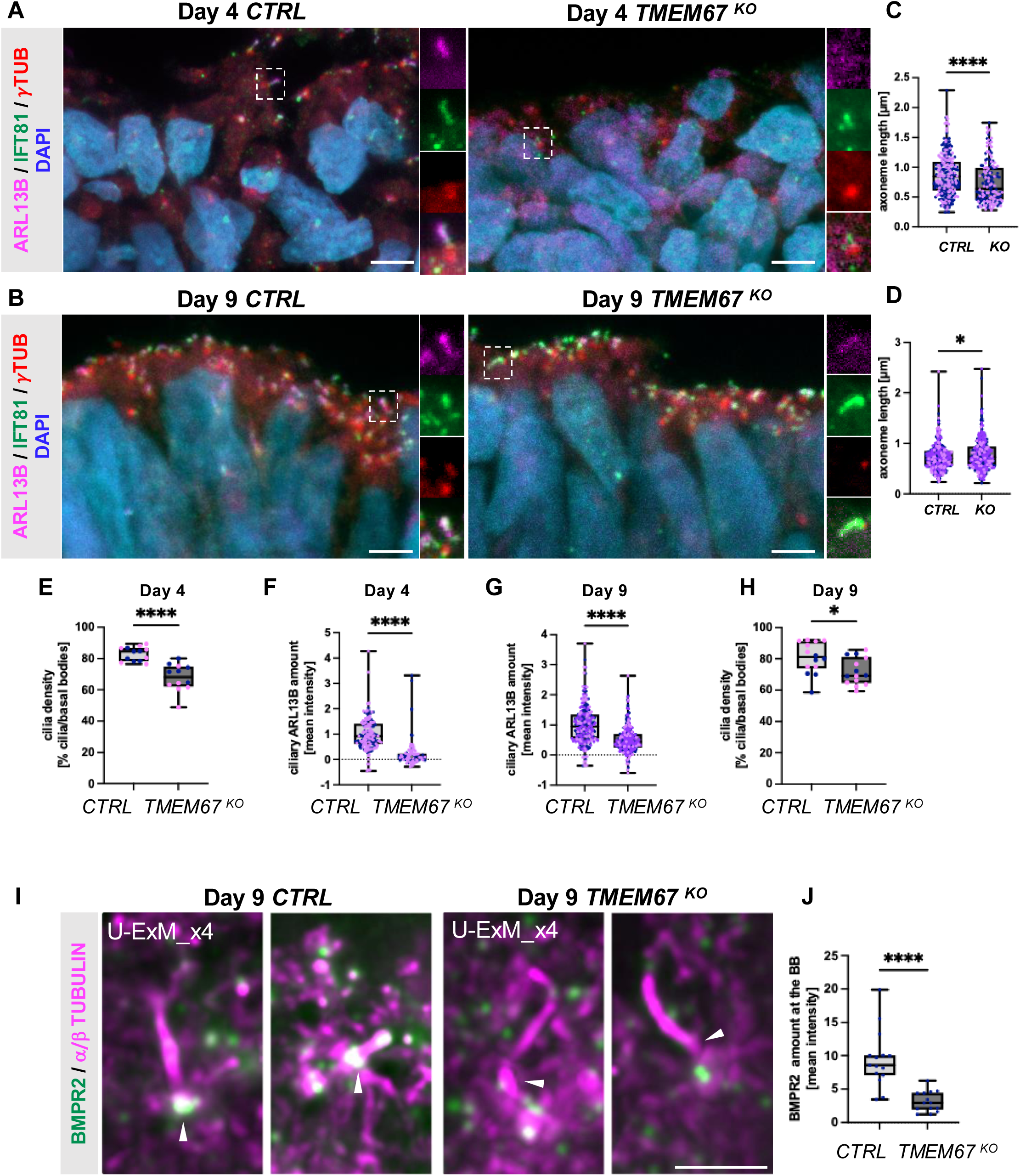
Altered ciliogenesis on TMEM67-deficient dorsal spinal progenitors coincides with defective BMPR2 recruitment at the ciliary base. **(A, B)** Immunofluorescence of cilia on *CTRL* and *TMEM67^KO^* spinal organoids at day 4 **(A)** and 9 **(B)** of differentiation. Cilia are labeled in green by IFT81 and magenta by ARL13B. Basal bodies are labeled in red by gTUBULIN. Magnified areas are indicated by white dotted rectangles and the corresponding magnified images of cilia are displayed on the right. The scale bar represents 5 µM. **(C, D)** Quantification of cilia length in on *CTRL* and *TMEM67^KO^* organoids at day 4 **(C)** and day 9 **(D)** of the differentiation. Data are shown as mean ± SEM. **(E-H)** Quantifications of cilia density and ciliary ARL13B amount in *CTRL* and *TMEM67^KO^* organoids at day 4 **(E, F)** and day 9 **(G, H)** of the differentiation. Data are shown as mean ± SEM. Each dot’s color represents a distinct experimental replicate (N≥2). Asterisks denote statistical significance according to unpaired t tests with Welch’s correction (*P < 0.05, ****P < 0.0001). **(I)** Immunofluorescence images of cilia and BMPR2 on *CTRL* and *TMEM67^KO^* spinal organoids at day 9 of differentiation obtained by Ultra Expansion Microscopy. White arrowheads point to basal bodies. BMPR2 receptor localizes at the basal body in cilia from *CTRL* organoids but is reduced at basal bodies of cilia in *TMEM67 ^KO^* organoids. **(J)** Quantification of BMPR2 amount at the basal bodies (BB) in on *CTRL* and *TMEM67^KO^* organoids at day 9 of the differentiation. Data are shown as mean ± SEM (N=1). Asterisks denote statistical significance according to unpaired t tests with Welch’s correction (****P < 0.0001). Scale bar, 5 µm.

## DISCUSSION

### Primary cilia as direct regulators of human dorsal spinal cord patterning

In this study, we uncover a direct and previously unappreciated role for the ciliopathy protein TMEM67 in the patterning of the human dorsal spinal cord. Using human dorsal spinal cord organoids, we show that loss of TMEM67 selectively disrupts specification of the roof plate and the dp1/dI1 lineage, while sparing—or even expanding—more intermediate dorsal identities. These patterning defects are accompanied by abnormal ciliary morphology and a highly penetrant loss of ARL13B localization to primary cilia, indicating a functional impairment of ciliary signaling.

A role for primary cilia in dorsal spinal cord patterning has previously been described in mouse models carrying mutations in ciliary genes such as *Arl13b*, *Fkbp8*, or *Tulp3*. In these studies, dorsal defects were largely interpreted as indirect consequences of ectopic Hedgehog (HH) pathway activation^25, 26, 27^. Notably, the phenotype observed in our human organoids closely resembles that of Arl13b mutant mice, which exhibit reduced *Gdf7* expression in the roof plate and altered specification of the dp1/dI1 (Atoh1⁺) lineage, despite preserved Wnt pathway activity. Importantly, in mice this phenotype was absent when *Arl13b* was conditionally deleted in dorsal progenitors using a Pax3-Cre driver, leading to the conclusion that dorsal patterning defects arose non–cell-autonomously from ventral tissues with aberrant Shh signaling and altered Gli activator-to-repressor balance^25^. Although we cannot exclude the possibility that a shift in GLI activator/repressor balance might antagonize BMP-mediated dorsal induction and participate in *TMEM67^KO^* organoids phenotype. However, since we only observe a slight increase in *GLI1* and *PTCH1* expression at day 9 in mutant organoids and predominantly generate PAX3-positive dorsal progenitors, we think our findings argue against such an indirect mechanism in the human context (Supplementary Figure 3G–K).

TMEM67 has also been implicated in WNT pathway regulation in other developmental and cellular contexts^42, 39^. However, our transcriptomic and functional analyses indicate that WNT signaling is not detectably altered in TMEM67-deficient dorsal progenitors and therefore cannot account for the early patterning defects we observe. Most interestingly, dorsal patterning abnormalities in *Arl13b* mutant mice were restricted to the posterior hindlimb level of the spinal cord^26^. This raises the intriguing possibility that anteroposterior identity modulates the sensitivity of dorsal progenitors to ciliary dysfunction. Future experiments manipulating rostro-caudal identity in spinal organoids may help clarify how regional differences in cilia composition or stability influence dorsal patterning outcomes.

### Cilia-dependent modulation of BMP signaling in human neural progenitors

BMP signaling is a central driver of dorsal spinal cord patterning, controlling both roof plate formation and the specification of dorsal progenitor domains. While BMP4 stimulation robustly induces SMAD1/5/9 phosphorylation and BMP target gene expression in control organoids, this response is significantly attenuated in TMEM67-deficient organoids. Notably, the spatial restriction of SMAD activation to the outer layers of spinal organoids—corresponding to dorsal progenitors exposed to higher BMP concentrations—is preserved in *TMEM67* mutants, although signal intensity is reduced (Figure 3A-C). These findings support a model in which primary cilia do not define the spatial range of BMP signaling, but instead modulate signal amplitude or timing, thereby influencing threshold-dependent fate decisions such as dp1 specification. Importantly, the reduced BMP response cannot be attributed to decreased expression of BMP ligands, receptors, or SMAD effectors, nor to gross defects in SMAD nuclear localization (Figure 3A, D-G). Instead, our data point to a role for primary cilia in early BMP signal transduction. We show that BMP4 stimulation induces recruitment of the type II BMP receptor BMPR2 to the base of the primary cilium in dorsal spinal progenitors, and that this recruitment is impaired in the absence of TMEM67 (Figure 4I, J). Given the established function of the ciliary transition zone as a selective gate controlling protein entry and exit, our findings suggest that TMEM67 facilitates BMP signaling by enabling efficient ciliary localization of BMPR2. Several components of the BMP and TGFβ pathways have been reported to localize to primary cilia in human (^43^ and Felix Hoffmann unpublished data), and mouse they were shown to mediate BMP signaling in the osteogenic lineage (^44,45^). It will be important in future work to confirm BMP or TGFβ pathway components localization to cilia in human spinal progenitors and how their trafficking to cilia indeed regulate their function.

### Implications for Joubert syndrome and human neurodevelopment

Temporal analysis of TMEM67^KO^ organoids reveals that dorsal patterning defects arise early at the progenitor stage. Markers of the dorsal-most progenitor domain dp1, in particular *ATOH1*, are consistently reduced, while transcriptional programs associated with intermediate dorsal progenitors are expanded. This imbalance is accompanied by downregulation of candidate regulators of the dp1/dI1 lineage, such as *ROBO3*, *CNTN2*, and *TLX3* (data not shown), suggesting a failure to activate the full dp1 gene regulatory network.

These early progenitor defects translate into a pronounced loss of dI1 interneurons at later stages, as indicated by reduced expression of *BARHL1*, *BARHL2*, *LHX2*, and *LHX9*, together with spatial disorganization of dorsal neuronal layers. Thus, impaired cilia-dependent BMP signaling has lasting consequences for dorsal spinal cord circuit assembly.

Spinal cord abnormalities have received relatively little attention in JBTS, yet histological analyses of patient samples have reported defects in the dorsal horns^29, 30^. Our findings provide a plausible developmental mechanism linking JBTS-associated transition zone defects to impaired BMP-dependent dorsal interneuron specification. Given the role of dI1 neurons in proprioceptive and sensory relay circuits, subtle alterations in their development may contribute to the motor and sensory deficits characteristic of ciliopathies.

Notably, the conservation of BMP signaling defects in RPGRIP1L-deficient organoids (data not shown) suggests that dysregulation of cilia-dependent BMP signaling may represent a shared pathogenic mechanism across multiple JBTS genes. This raises the possibility that targeted modulation of BMP pathway activity could be explored as a therapeutic strategy in specific ciliopathy contexts.

### Limitations and future perspectives

While spinal organoids provide a powerful platform to study early human neural development, they do not fully recapitulate the complex morphogen gradients and tissue interactions present *in vivo*. Future studies using additional iPSC as well as patient-derived iPSC clones, but also more complex systems such as dorso-ventral spinal assembloids may help refine BMP gradient dynamics and further dissect cilia-dependent signaling mechanisms. In addition, single-cell transcriptomic analyses and rescue experiments restoring TMEM67 or targeting BMPR2 ciliary localization will be essential to validate and extend our model.

## MATERIAL AND METHODS

### Ethics

The work performed in this manuscript complies with all relevant ethical regulations. The obtaining and manipulation of human iPSC lines were performed under ethical agreement DC- 2025-7003 from the French Ministry of Higher Education and Research and GMO authorization number DUO-9509.

### Generation and amplification of *TMEM67^KO^* and Control mutant hiPSC lines

Characterization and generation of TMEM67-deficient hiPSC clones from the PCli033-A cell line were already described^33^. Briefly, TMEM67-deficient hiPSCs and their isogenic controls in the Pheno2 cell line were obtained using CRISPR/Cas9 gene editing combining 2 guide RNAs targeting exon 1 and exon 27 of *TMEM67* to generate a large deletion of TMEM67 (Δexon1-exon27). One homozygous *TMEM67^KO^* clone and one heterozygous control clone were used for further analyses. For all clones, genomic stability was assessed by detection of recurrent genetic abnormalities using the iCS-digitalTM PSC test, provided as a service by Stem Genomics (https://www.stemgenomics.com).hiPSCs were thawed in presence of Rock-inhibitor Y-27632 (5µM, Stemgent-Ozyme #04-0012) and cultured under standard conditions at 37°C in mTeSR+ medium (Stem Cell Technologies #100-0276) on Matrigel (Corning, VWR #354277) coated plates upon confluency of 70-80%. Passaging was performed using ReLeSR (Stem Cell Technologies #05872) and testing for potential mycoplasma contamination was performed regularly by Eurofins genomic (Mycoplasmacheck). Accutase (Stem Cell Technologies #07920) was used for the dissociation of hiPSC colonies into single cells.

### Differentiation of hiPSCs into dorsal spinal organoids

Differentiation of control and mutant hiPSCs was performed as schematized in Figure 1A. After amplification, hiPSC lines were dissociated into single cells using Accutase (Stem Cell Technologies #07920) and resuspended in differentiation medium N2B27 [vol:vol; Advanced DMEM/F-12 (Gibco) and Neurobasal Medium (Gibco)] supplemented with N2 (Thermo Fisher #17502048), B27 without Vitamin A (Thermo Fisher #12587010), penicillin/streptomycin 1 % (Thermo Fisher #15140122), β-mercaptoethanol 0.1 % (Thermo Fisher #31350010). Cells were seeded in ultra-low attachment dishes (Corning #3261) to allow EB formation. The N2B27 differentiation medium was used throughout the whole differentiation process, supplemented with specific small molecules and recombinant proteins at different time points as follows: Rock-inhibitor Y-27632 (5 μM; Stemgent-Ozyme #04-0012) was added from day 0 to day 2, CHIR-99021 (3 µM; Stemgent-Ozyme #04-0004) from day 0 to day 4, SB431542 (20 μM; Stemgent-Ozyme #04-0010) from day 0 to day 3 and LDN 193189 (0.1 μM; Stemgent-Ozyme #04-0074) from day 0 to day 4. BMP4 recombinant protein (5ng/ml) was added from day 4 to 5 and Retinoic Acid (100 nM; Sigma #R2625) from day 4 to day 9. γ-Secretase inhibitor DAPT (10 µM; Tocris Bioscience #2634) was added from day 9 to the end of differentiation. Medium was changed every other day.

### EB embedding and Cryosectioning

Spinal organoids were collected at different time points during differentiation, rinsed with PBS and fixed in cold PFA (4 %) for 7-12 min at 4 °C. EBs were rinsed in PBS and incubated in 30 % sucrose in PBS until completely saturated. Cryoprotected EBs were embedded in OCT embedding matrix (Cell Path #KMA-0100-00A) and stored at -80 °C. 12 µM cryostat sections were prepared and stored at -80 °C.

### Immunofluorescence

Cryostat sections of organoids were washed in PBS/0.1% Triton X-100. Optional steps for permeabilization with PBS/0.5 %Triton X-100 for 10 min, post-fixation with MeOH (100 %) for 10 min or antigen-retrieval with Citrate-Buffer were performed depending on the antibody and specimen. After washing, cryosections were incubated with a blocking solution containing 10 % NGS in PBS/0.1% Triton-X100 for at least 2 hours. The sections were incubated with primary antibodies (Table S2) diluted in PBS/0.1 % Triton-X100 and 1 % NGS overnight at 4 °C. Next day, the slides were washed 3 x 5 min with PBS/0.1 % Triton-X100 and incubated with the secondary antibodies (Table S2) diluted in PBS/0.1 % Triton X-100 and 1 % NGS for 2 hours. After final washing steps with PBS/0.1 %Triton X-100 for at least 1 hour, sections were mounted with Vecta-Shield (Vector #H-1000) or Mowiol (Roth #0713.2) optionally containing DAPI (Merck #1.24653).

### Ultrastructure-Expansion Microscopy (U-ExM)

U-ExM was performed on fixed sections of spinal organoids according to a protocol adapted from Gambarotto et al. 2019. Briefly, frozen sections on slides were thawed at RT for 2 min. A double-sided sticky spacer of 0.3-mm thickness (IS317, SunJin Lab Co.) was stuck on the slide around the section of interest. The crosslinking prevention step was performed by incubation in 2% acrylamide (AA; A4058, Sigma-Aldrich) and 1.4% formaldehyde (FA; F8775, Sigma-Aldrich) in a total volume of 300 µl at 37°C for 3 h. After removing the first solution, the gelation step was done by adding 130 µl of monomer solution composed of 75 μl of sodium acrylate [stock solution at 38% (w/w) diluted with nuclease-free water, 408220, Sigma-Aldrich], 37.5 μl of AA, 7.5 μl of N,N′-methylenbisacrylamide (BIS, 2%, M1533, Sigma-Aldrich), 10 μl of 10× PBS together with ammonium persulfate (APS, 17874, Thermo Fisher Scientific) and tetramethylethylenediamine (TEMED, 17919, Thermo Fisher Scientific) at a final concentration of 0.5%, for 1 h and 30 min at 37°C. Of note, APS and TEMED have to be added at the last minute to avoid premature polymerization. A 24-mm coverslip was added on top to close the chamber. Next, the coverslip and the spacer were removed, and the slide was submerged in a 50 ml Falcon tube filled with denaturation buffer pre-heated at 95°C [200 mM SDS, 200 mM NaCl, 50 mM Tris base in water (pH 9)]. The Falcon tube was then incubated for 2 h at 95°C in a water bath. Finally, the gel was rinsed and expanded in three successive ddH2O baths, before processing to the immunostaining.

### Image acquisition

Fluorescence image acquisition of patterning and differentiation markers was performed at room temperature using a Zeiss Observer Z1 equipped with an Apotome module, an Axiocam 506 monochrome cooled CCD camera and a 40x oil objective with a NA 1.3. All images were processed with Zen software (blue edition). Image analyses and figure preparation were performed using Fiji (ImageJ; National Institutes of Health). For cilia analyses, fluorescence images were acquired at room temperature using an inverted confocal microscope (LSM 980, Carl Zeiss AG), a 63x oil objective with a NA 1.4, and a monochrome charge-coupled device camera. Z-stacks with 0.2 µM steps were acquired. Images were processed with Zen software (black edition). Image analyses and figure preparation were performed using Fiji (ImageJ; National Institutes of Health).

### Quantification of fluorescent staining and statistical analysis

Fluorescent staining and protein band intensities were quantified using Fiji (ImageJ; National Institutes of Health). Intensity measurements of ciliary proteins were performed on unprocessed images (raw data) as described before ^33^. After quantifications were performed, representative images were processed by means of background subtraction and contrast settings via Adobe Photoshop CS2.

For quantitative analyses of nuclear staining, automated nuclear segmentation was first performed on the DAPI channel using Cellpose 2.0. Intensity of the staining was measured for each nucleus by uploading the corresponding ROI mask into Fiji (ImageJ; NIH) on the corresponding MAX-projection image. Mean intensity was computed for each ROI, and a threshold was used to differentiate marker-positive cells from background immunostaining. Only nuclei exceeding this threshold were included in the calculation of the positive cell ratio. At least 10 images per clone, from 2 independent experiments of differentiation were used for area quantification.

All data are presented as mean ± SEM. Two-tailed t test with Welch’s correction was performed for all data in which two datasets were compared. Two-tailed *t* test Analysis of variance (ANOVA) and Tukey honest significance difference (HSD) tests were used for all data in which more than two datasets were compared. Following statistical significances were considered: \**P* < 0.05, \*\**P* < 0.01, \*\*\**P* < 0.001 and \*\*\*\**P* < 0.0001. All statistical data analysis and graph illustrations were performed using GraphPad Prism (GraphPad Software).

### RNA isolation and qRT-PCR analyses

hiPSC-derived organoids from 3 independent experiments were collected at day 4, 9 and 14 of the differentiation. Material from one control (*TMEM67*^+/-^) and one mutant (*TMEM67*^-/-^) clone of the Ph2 hiPSC line was collected. RNAs were extracted using the RNeasy Kit (Qiagen #74104) and RNAse-free DNase Set (Qiagen #79254) and retro-transcribed into cDNA using the Verso™ RT-PCR Kit (ThermoFisher # AB-1453/A). For quantitative real-time PCR, 50 ng of cDNA of each sample was used in a Maxima SYBR Green/ROX qPCR Master Mix 2x (Thermo Scientific #K0222). Triplicate reactions for each sample were prepared and the samples were run in a Step One Real-Time PCR System Thermal Cycling Block (Applied Biosystems #4376357). Primer pairs are listed in Table S1. The analysis of real-time data was performed using the included StepOne Software version 2.0 and graphs were generated using GraphPad Prism (GraphPad Software).

### Bulk RNAseq analysis on human spinal organoids

After organoids collection and RNAs extraction as detailed in the preceding paragraph, RNA quality was determined by measuring the RNA integrity number (RIN) via the High Sensitive RNA Screen Tape Analysis kit (Agilent Technologies #5067) on the TapeStation system (Agilent Technologies). A RIN above 9.5 was considered as good quality RNA and 250 ng RNA in a total volume of 25 µl was prepared per sample for further procedure. Bulk RNAseq was realized by the Genotyping and Sequencing Core Facility of the Paris Brain Institute (iGenSeq, ICM Paris). RNAseq data processing was performed in Galaxy under supervision of the InforBio platform (IBPS, Paris). Paired-end RNA reads were aligned against the *homo sapiens hg38* genome by using the STAR aligner (v2.7.10, Galaxy wrapper v4)^46^ and the gene annotations gtf files GRCm39.109 and GRCh38.109, respectively. Quality of sequencing was controlled with fastQC (Galaxy wrapper v0.74+galaxy0) and MultiQC (Galaxy wrapper v1.11+galaxy1)^47^. Gene expression was assessed by counting sequencing reads aligned to gene exons with featureCounts (Galaxy wrapper v2.0.3+galaxy2)^48^. Raw counts were further converted to normalized gene expression values using the log2(tpm+1) transformation where tpm is the count of transcript aligned reads per length of transcript (kb) per million of total mapped reads (Galaxy tool "cpm_tpm_rpk" v0.5.2). Principal Component Analyses (PCA) were performed using r-factominer (v.2.9, Galaxy tool id "high_dimensions_visualisation" v4.3+galaxy0) and heatmap visualizations were produced with the Galaxy tool "high_dim_heatmap" (v3.1.3+galaxy0) using the normalized gene expression values. Differentially expressed genes (DEGs) were selected from the gene raw read counts using DESeq2 (Galaxy wrapper v2.11.40.8+galaxy0)^49^ and the Benjamini-Hochberg p-adjusted cutoff 0.01. DESeq2 statistical tables were used for generation of Volcano Plots (Galaxy tool id "volcanoplot" v0.0.5)^50^.

## Acknowledgement

Image acquisition was carried out at the IBPS Imaging Facility. The IBPS Imaging facility is supported by Region-Île-de-France, Sorbonne-University and CNRS. We thank the iGenSeq sequencing platform at the ICM for reliable sequencing and fast processing of our samples. We are very grateful to Vanessa Ribes and Pascal Gilardi Hebenstreit for providing us with protocols and help in culture and differentiation approaches. We acknowledge the worldwide contributions of users and developers to the Galaxy project (https://galaxyproject.org/) and all the upstream authors and contributors of the software ecosystem we use.

We further thank Felix Hoffmann (University Hospital Tübingen, Germany) to share data on BMP pathway components and cilia.

This work was supported by funding to SSM from the *Fondation pour la Recherche Médicale* (Equipe FRM *EQU202503020010*), from the *Fondation pour la Recherche sur le Cancer* (ARC PJA 2024080008603), from the *Agence Nationale de la Recherche* - ERA-NET Neuron 2021 “NDCil” (ANR-21-NEU2-0009-03) project and PRC AAPG2024 “CiCerO” project, from the *Fondation Maladies rares* and the *Association Mieux vivre avec le syndrome de Joubert* under the reference “ASSOJoubert_2024_Schneider”. AW received funding from the *German Research Foundation* (DFG; WI 5451/1-1).

